# Use of Epivolve phage display to generate a monoclonal antibody with opsonic activity directed against a subdominant epitope on extracellular loop 4 of *Treponema pallidum* BamA (TP0326)

**DOI:** 10.1101/2023.05.13.540667

**Authors:** Mary Ferguson, Kristina N. Delgado, Shannon McBride, Isabel C. Orbe, Carson J. La Vake, Melissa J. Caimano, Qiana Mendez, Trevor F. Moraes, Anthony B. Schryvers, M. Anthony Moody, Justin D. Radolf, Michael Weiner, Kelly L. Hawley

**Affiliations:** Department of Molecular Sciences, Abbratech, Branford Connecticut, USA; Department of Medicine, UConn Health, Farmington, CT USA; Research and Development Abcam, Branford Connecticut, USA; Department of Pediatrics, UConn Health, Farmington, CT USA; Department of Molecular Biology and Biophysics, UConn Health, Farmington, CT USA; Department of Biochemistry, University of Toronto, Canada; Department of Microbiology, Immunology and Infectious Diseases, University of Calgary, Calgary, Canada; Duke Human Vaccine Institute, Durham, NC USA; Department of Pediatrics, Duke University Medical Center, Durham, NC USA; Department of Integrative Immunology, Duke University Medical Center, Durham, NC USA; Department of Immunology, UConn Health, Farmington, CT USA; Department of Genetics and Genome Sciences, UConn Health, Farmington, CT USA; Division of Infectious Diseases and Immunology, Connecticut Children’s, Hartford, CT USA

**Keywords:** Syphilis, *Treponema pallidum*, outer membrane protein, BamA ECL4, opsonic antibody, monoclonal antibody, subdominant epitope, *Pyrococcus furiosus* thioredoxin.

## Abstract

Syphilis, a sexually transmitted infection caused by the spirochete *Treponema pallidum* (*Tp*), is resurging globally. Opsonic antibodies (Abs) targeting surface-exposed epitopes of the spirochete’s outer membrane proteins (OMPs) are believed to promote macrophage-mediated clearance of the bacterium during infection and are presumed to be key to vaccine development. *Tp*’s repertoire of outer membrane proteins includes BamA (β-<u>b</u>arrel <u>a</u>ssembly <u>m</u>achinery subunit <u>A</u>/TP0326), the central component of the molecular machine that inserts newly exported OMP precursors into the OM lipid bilayer. BamA is a bipartite protein consisting of an 18-stranded β-barrel with nine extracellular loops (ECLs) and five periplasmic POTRA (<u>po</u>lypeptide <u>tr</u>ansport-<u>a</u>ssociated) domains. Antisera directed against BamA ECL4 promote internalization of *Tp* by rabbit peritoneal macrophages. Herein, we employed a novel two-stage, phage display strategy, termed “Epivolve” (for <u>epi</u>tope <u>evol</u>ution), to generate five site-directed murine monoclonal Abs (mAbs) targeting a centrally located peptide (S2) of BamA ECL4. Each of the five mAbs demonstrated reactivity by immunoblotting and ELISA to nanogram amounts of BamA ECL4 displayed by a *Pyrococcus furiosus* thioredoxin (*Pf*Trx) scaffold (*Pf*Trx^BamA/ECL4^). One mAb containing a unique amino acid sequence in both light and heavy chains showed activity in an opsonophagocytosis assay employing murine bone marrow-derived macrophages. Mice and rabbits hyperimmunized with *Pf*Trx^BamA/ECL4^ produced opsonic antisera that strongly recognized the ECL presented in a heterologous scaffold and overlapping ECL4 peptides including S2. In contrast, Abs generated during *Tp* infection of mice and rabbits poorly recognized the peptides, indicating that S2 contains a subdominant epitope. Epivolve, which circumvents the natural immune response, can be utilized for the generation of mAbs that target subdominant opsonic epitopes in ECLs of *Tp* OMPs.

## INTRODUCTION

Syphilis is a multistage, sexually transmitted infection caused by the highly invasive and immunoevasive spirochete *Treponema pallidum* subspecies *pallidum* (*Tp*) (1, 2). Since the start of the new millennium, syphilis has undergone a dramatic resurgence in the United States, particularly among men who have sex with men (3) in addition to posing an ongoing threat to at-risk populations in resource-poor nations (4). These alarming trends underscore the urgent need for new control strategies, including vaccines (4). It is generally believed that an improved understanding of host defenses responsible for spirochete clearance mechanisms is essential for syphilis vaccine design. The appearance of opsonic antibodies (Abs) directed against an increasingly broad spectrum of surface-exposed antigens as infection proceeds presumably tips the balance in favor of the host during its protracted battle with the ‘stealth pathogen’ (5). The principal targets of these opsonic Abs are believed to be the extracellular loops (ECLs) of the spirochete’s rare outer membrane proteins (OMPs) (6). *Tp*’s repertoire of OMPs includes BamA (β-<u>b</u>arrel <u>a</u>ssembly <u>m</u>achinery subunit <u>A;</u> TP0326), the central component of the molecular machine that inserts newly exported OMP precursors into the OM lipid bilayer (6, 7, 8). *Tp* BamA is a bipartite protein consisting of an 18-stranded β-barrel with nine ECLs and a periplasmic arm containing five POTRA (<u>po</u>lypeptide <u>tr</u>ansport-<u>a</u>ssociated) domains (9). We previously reported that antisera directed against BamA ECL4 promote opsonophagocytosis of *Tp* by rabbit macrophages (10). These results suggested that ECL4 Abs generated during infection contribute to spirochete clearance and that *Tp* BamA ECL4 might serve as a prototype for potentially protective Ab-ECL interactions.

Monoclonal Abs (mAbs) are powerful tools for identifying new vaccine antigens and defining natural and conformationally specific protective epitopes (11). While mAbs have been used extensively to study protective epitopes for viral infections (12, 13, 14), only a handful of studies have utilized mAbs for vaccine development against bacterial pathogens (15). In the early 1980’s, mAbs were generated against a number of *Tp* immunogens (16, 17, 18); however, it subsequently was determined that the targets of these mAbs are subsurface lipoproteins(19, 20). Enhanced 3D modeling of *Tp*’s repertoire of OMPs (6) – the *Tp* ‘OMPeome’ – now makes possible the use of mAb technologies to study protective immunity in syphilis at the structural and molecular level. Herein, we employed a novel, two-stage, phage display strategy, termed “Epivolve” (for <u>epi</u>tope <u>evol</u>ution; **Figure 1**), to generate a site-directed murine mAb with opsonic activity directed against a subdominant epitope on ECL4 of *Tp* BamA. We found that Abs against this epitope are often absent in syphilitic sera but can be generated by hyperimmunization with the ECL displayed on a protein scaffold.

**Figure 1.**
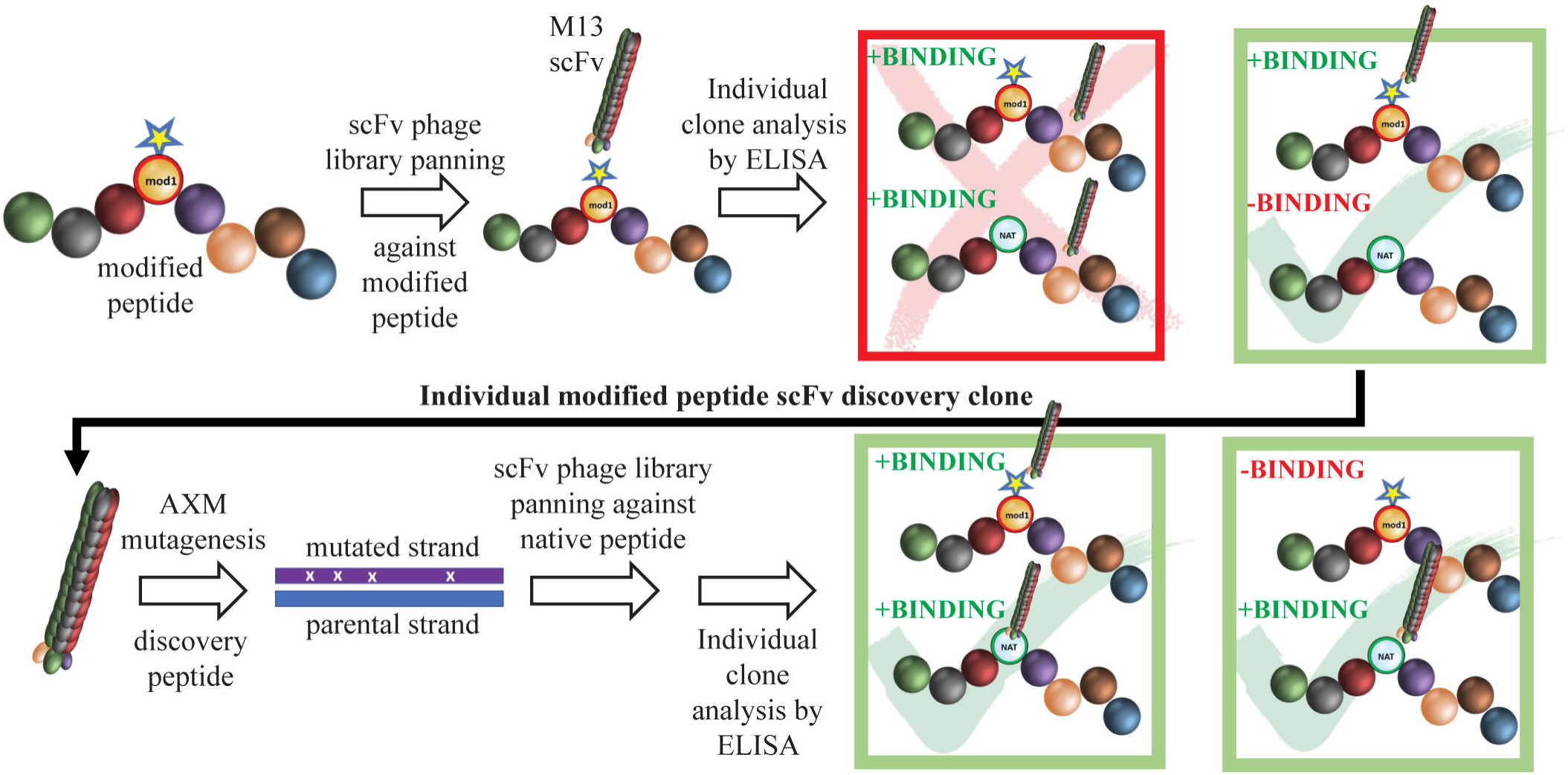
Schematic of mAb generation using Epivolve. A peptide incorporating a non-native amino acid at a desired site is used to pan a scFv phage library for peptide binders. Phages that bind the modified, but not the native, peptide undergo AXM mutagenesis (22) to generate phages that recognize the native peptide +/-the modified peptide with high affinity.

## MATERIALS AND METHODS

### Ethics statement

Animal experimentation was conducted following the *Guide for the Care and Use of Laboratory Animals* (8th Edition) in accordance with protocols reviewed and approved by the UConn Health Institutional Animal Care and Use Committee under the auspices of Animal Welfare Assurance A3471-01.

### Bacterial strains and plasmids

The *Escherichia coli* strains TG1 and AXE688 (21, 22, 23) were purchased from Lucigen Corporation (Middleton, WI). *E. coli* NEB^®^ 5-alpha and NEBExpress^®^ strains were purchased from New England BioLabs (Ipswich, MA). The template plasmid for all phage display libraries is a derivative of the phagemid pIT2 (24) with a human single-chain variable fragment Ab (scFv) fused to the coat protein III of bacteriophage M13, constructed at and kindly supplied by AxioMx Inc., an Abcam Company (“Abcam”).

### Phage library construction

Novel pre-defined complementarity determining regions (PDC) libraries have been described previously (25) (see Figure S1). Briefly, several thousand short oligos were synthesized for each of Ab complementarity determining regions (CDRs) HC1, HC2, LC1 and LC2. The specific sequences were chosen from successful phage display screens against over 1000 different antigens, including peptides and proteins. Furthermore, the chosen CDRs were from scFvs that expressed protein at high levels in *E. coli*. CDR HC3 and LC3 sequences were synthesized with varying lengths using an NNK codon. The potential diversity of the library was over 10^18^ of which we sampled through ∼10^12^ for this study.

### BamA ECL4 Epivolve peptides

The homology model of *<u>Tp</u>* BamA (**Figure 2A**) was previously generated (PDB is downloadable from https://drive.google.com/file/d/1EurEnlwAiqtsUm8t-jC3Xuz5e7nV45mT/view?usp=sharing&export=download) (6). The predicted B-cell epitopes (BCEs) were identified in ElliPro (26) using a threshold setting of 0.8. BamA ECL4 was divided into three overlapping peptides: S1 (residues 567-583; VIRVNGGVDFRVVKNFY); S2 (residues 577-594; RVVKNFYDKDNNQPFDL); and S3 (residues 584-602; DKDNNQPFDLTVKEQLNWT). Each peptide contains a centrally located aspartic acid residue that served as the modified site (**Figure 2B**). Native and modified BamA ECL4 peptides and an irrelevant peptide, all with N-terminal biotin, were purchased from Biopeptek Pharmaceuticals, LLC (Malvern, PA).

**Figure 2.**
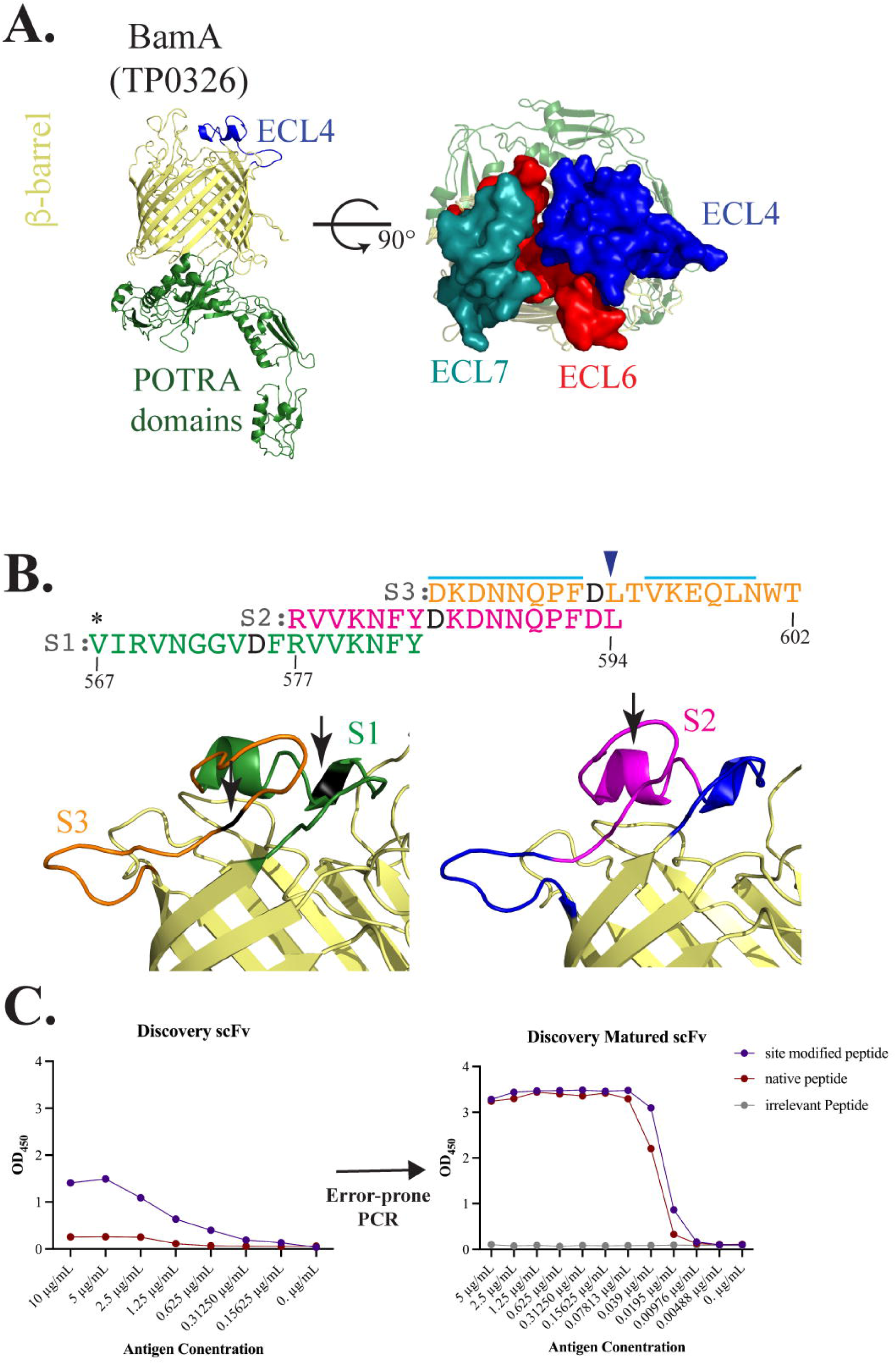
BamA ECL4 is a target antigen for mAb generation. (**A**) Ribbon diagram for the structural model of *Tp* BamA (TP0326) depicting the β-barrel, ECL4 and the five periplasmic polypeptide-transport-associated (POTRA) domains. ECLs 4, 6 and 7 form a dome that occludes the barrel opening. (**B**) Sequences of the three overlapping ECL4 peptides (S1-S3) used for Epivolve. The modified aspartic acid residue in each peptide is represented in black. A light blue line indicates predicted linear B cell epitopes in S2 and S3. Asterisk indicates an additional residue added to centrally position the modified residue in the S1 peptide. Blue arrow above the sequence indicates the glutamine to leucine substitution in the Mexico A strain (10). (**C**) Representative titration ELISAs using native and modified peptides, pre-and post-error-prone PCR.

### Epivolve discovery phase phage display screen

Immunoplates (Nunc Maxisorp) were coated with NeutrAvidin (ThermoFisher) overnight (ON) at 4 °C. Plates were washed with PBS and then blocked with 2% nonfat dry milk in PBS (MPBS). After a PBS wash, plates were coated for 1 h with biotinylated peptide (10 µg/ml). After a PBS wash and block with MPBS, phage library was added at 1 x 10^12^ phage/ml and incubated for 1 h at room temperature (RT). After rigorous washing with PBS containing 0.1% Tween 20 (PBST), bound phage were recovered by addition of trypsin (100 µl/well), transduced into exponentially growing *E. coli* TG1 (Lucigen Middleton, WI) for 30 min at 37 °C, and then grown overnight in 2YT containing ampicillin (100 µg/ml) and 1% glucose at 30 °C. The following day, cultures were diluted into fresh medium of 2YT containing ampicillin (100 µg/ml) and 1% glucose and incubated at 37 °C with shaking until OD_600_ = 0.4. KM13 helper phage were added at a multiplicity of infection (MOI) of 10:1 and incubated at 37 °C for 30 min. Transduced cells were then pelleted and incubated overnight in 2YT containing ampicillin (100 µg/ml) and kanamycin (50 µg/ml). The resulting phage supernatant was applied to another antigen coated immunoplate and the entire process was repeated for a total of three rounds. The corresponding nonbiotinylated and nonphosphorylated peptides were added as competing antigens during the second and third rounds of panning to remove scFv-phage molecules that preferentially bound to the modified neoepitope. After the third round, supernatants from 88 single scFv-containing colonies was tested by ELISA for binding against the respective modified and native peptides; NeutrAvidin alone was used as a negative control.

### Affinity maturation using AXM mutagenesis

Mutagenized libraries for directed evolution were previously generated utilizing thiol-protection of one of a pair of common PCR primers (21) (Figure S2). The coding region for the selected Ab was amplified under error prone PCR as previously described (22) using 0.5 mM manganese chloride to facilitate mutagenic nucleotide incorporation. The reverse primer containing phosphorothioate linkages on its 5′ end. The resulting double-stranded DNA was treated with T7 exonuclease (New England Biolabs, Ipswich, MA) to selectively degrade the unmodified strand of the dsDNA molecule. The resulting single-stranded DNA, or ‘megaprimer’, was then annealed to the uracilated, single-stranded circular phagemid DNA and used to prime *in vitro* synthesis by DNA polymerase (New England Biolabs). The ligated, heteroduplex product was then transformed into *E. coli* TG1 cells (Lucigen), where the uracilated strand is cleaved *in vivo* by uracil N-glycosylase, favoring survival of the newly synthesized, recombinant strand containing the megaprimer (21, 22).

### Epivolve maturation phase phage display screen

The template phagemid for second generation affinity maturation libraries were based on evolved scFv sequences identified from phage display panning of the above mutagenized libraries. Affinity maturation phage libraries were generated as previously described for AXM mutagenesis (21, 22) using AXE688 electrocompetent cells and optimized conditions for AXL40 and AXL41 template phagemids. The mutagenized libraries on average have an estimated average diversity of 10^7^.

### IgG production

Heavy and light chain sequences from successful scFv clones were identified *via* Sanger sequencing (GeneWiz, South Plainfield, NJ). Heavy and light chain DNA were synthesized and cloned into a mouse pTT5 expression vector by BioBasic Inc. Plasmids were then transfected into human embryonic kidney cells (HEK293-E, National Research Council, Canada) using 293Fectin (Thermofisher). Six days post-transfection, IgG from the harvested supernatant was purified using a Protein A column (Cytiva Life Sciences, Marlborough, MA) and dialyzed to resuspension in PBS.

### Complementarity-determining regions (CDR) analysis

The heavy and light chain sequence for each mAb was submitted to the AbYsis website (http://www.abysis.org/abysis/sequence_input/key_annotation/key_annotation.cgi) to identify canonical class assignments for CDRs and unusual residues involved in antigen binding.

### Cloning of recombinant proteins

The *Pyrococcus furiosus* thioredoxin scaffold containing *Tp* BamA ECL4 (*Pf*Trx^BamA/ECL4^) containing N-terminal His-and C-terminal Avi-Tags and cloned into pET28a was described previously (27). The “empty” *Pf*Trx scaffold (*Pf*Trx^Empty^) was generated by self-ligating BamHI-digested *Pf*Trx^BamA/ECL4^. The *Pf*Trx^BamA/ECL4^ Mexico A construct was generated by reverse PCR using Q5 Hot Start High-Fidelity DNA polymerase (New England Biolabs, Inc.) in a 25 μl reaction containing 100 ng of *Pf*Trx^BamA/ECL4^ (Nichols) plasmid and *Pf*Trx^BamA/ECL4^ MexA–FW and –RV primers (see Table S1). The resulting amplicon was self-ligated using Kinase, Ligase & Dpnl Enzyme Mix (New England Biolabs, Inc.), transformed into Top10 cells and plated on LB containing kanamycin. Clones were confirmed by Sanger sequencing. Generation of codon-optimized *Tp* BamA β-barrel in pET23b was previously described (9). Codon-optimized C-lobe of *Neisseria meningitidis* transferrin binding protein B, named the ‘loopless’ C-lobe (TbpB-LCL) (28) and TbpB-LCL scaffold containing *Tp* BamA ECL4 (TbpB-LCL^BamA/ECL4^) were generated by gene synthesis (Azenta Life Sciences, Burlington, MA) and cloned into *Nde*I-*Xho*I-digested pET28a by In-Fusion cloning (Takara Bio USA, Inc., San Jose, CA), according to the manufacturer’s instructions. Oligonucleotide primers used in this study are presented in Table S1.

### Expression and purification of recombinant proteins

*Pf*Trx proteins were expressed in BirA-transformed *E. coli* BL21 (DE3) (BPS Bioscience, San Diego, CA) for *in vivo* biotinylation in Lysogeny Broth (LB) containing kanamycin (50 μg/ml), spectinomycin (50 μg/ml) and 50 μM D-biotin (Thermo Fisher Scientific, Waltham, MA) and then purified over Ni-NTA (Qiagen, Germantown, MD) as previously reported in (27). Soluble *Pf*Trx proteins were further purified by size-exclusion chromatography (SEC) over a Superdex 200 Increase 10/300 GL column (Cytiva, Marlborough, MA). Proteins used for immunization (see below) were dialyzed with PBS for 4 h at 4°C. Recombinant BamA β-barrel was expressed in *E. coli* C41 (DE3) cells, grown in LB containing 50 μg/ml kanamycin and the insoluble recombinant proteins were purified as previously described (10) and not subjected to SEC.

TbpB-LCL proteins were expressed in *E. coli* Gold (DE3) cells (Agilent) using Overnight Express Instant LB medium (Millipore Sigma, St. Louis, MO) containing 50 μg/ml kanamycin. Washed cell pellets were lysed in BugBuster (Novagen) containing lysozyme, DNAse and protease inhibitor cocktail. Following centrifugation, the soluble fraction was purified over Ni-NTA resin, washed once each with TbpB-LCL Wash Buffer A (50 mM Tris-HCl [pH 7.5], 500 mM NaCl, 10 mM imidazole) and Wash Buffer B (50 mM Tris-HCl [pH 7.5], 200 mM NaCl, 20 mM imidazole) and then eluted in Wash Buffer B containing 300 mM imidazole. Following elution, fractions containing TbpB-LCL proteins were further purified over a Superdex 200 Increase 10/300 GL column (Cytiva) in buffer containing 50 mM Tris-HCl (pH 7.5), 200 mM NaCl and 1 mM β-mercaptoethanol.

### Generation of antiserum in rat, mice and rabbits

Rat α-BamA ECL4 (BamA residues 568-602) antiserum was generated as described previously (10). For mouse polyclonal antisera, six-to eight-week-old C3H/HeJ mice (Jackson Laboratory) were primed by intradermal injections with 100 μl Freund’s Complete Adjuvant (1:1 v/v) containing 20 μg of *Pf*Trx^BamA/ECL4^ or *Pf*Trx^Empty^. Mice were boosted at 3, 5 and 7 weeks with the same volumes and amounts of protein in Freund’s Incomplete Adjuvant (1:1 v/v) and exsanguinated 9 weeks post-immunization. Serum from five mice was pooled, heat inactivated and then used in immunologic assays. Adult male New Zealand White (NZW) rabbits (Envigo, Indianapolis, IN) were primed by four subcutaneous injections and two intramuscular injections with 100 μl and 50 μl PBS-TiterMax (1:1 v/v) respectively, containing 200 μg of *Pf*Trx^BamA/ECL4^ or *Pf*Trx^Empty^. Rabbits were boosted at 3, 6 and 9 weeks with the same volumes and amounts of protein in PBS-TiterMax (1:1 v/v) and exsanguinated 12 weeks post-immunization.

### ELISA with murine IgG_2_ monoclonal antibodies

For titration ELISA, Maxisorp 96-well plates were coated with 50 μl/well of NeutrAvidin (ThermoFisher Scientific) at a final concentration of 1 μg/ml. The NeutrAvidin-coated plates were washed 3× with PBS and blocked with 1% BSA/PBS for 1 h at RT. The plates were then washed 3× with PBS and coated with 1 µg/ml biotinylated peptide antigen. Protein antigens were directly coated to the NeutrAvidin-free plate for 1 h. Plates were blocked with 1% BSA/PBS at RT for 1 h. Seven titrations (30, 7.5, 1.875, 0.47, 0.12, 0.007 and 0 μg/ml) of each Ab were diluted in 1% BSA/PBS, applied to the plates and incubated for 1.5 h. The plates were then washed 3× with PBST. AffiniPure goat α-mouse HRP (<u>h</u>orseradish <u>p</u>eroxidase; Jackson ImmunoResearch) was diluted 1:10,000 in 1% BSA/PBS, added to wells and then incubated for 1 h at RT. Plates were washed 3× with PBST. After the addition of Ultra TMB reagent (ThermoFisher Scientific), wells were developed for 5 min at RT and then stopped with 0.16 M H_2_SO_4_ (50 μl/well). ELISA signal (absorbance at 450 nm) was measured using an Envision plate reader (BD, East Rutherford, NJ).

### ELISA with syphilitic sera and BamA ECL4 antisera

ELISAs were conducted as previously described (27) with the exception of using biotinylated *Pf*Trx^BamA/ECL4^ and *Pf*Trx^Empty^ proteins added at 200 ng/well in PBS buffer containing 15% goat serum, 0.005% Tween 20 and 0.05% sodium azide followed by 1 h of incubation at RT. The optical density (450 nm) readings of serial dilutions for *Pf*Trx^BamA/ECL4^ were used to calculate area under the curve (AUC). The AUC for *Pf*Trx^Empty^ was subtracted from the AUC of each *Pf*Trx construct.

### Immunoblot analysis

To assess the reactivity of *Pf*Trx^BamA/ECL4^ construct with each mAb, a gradient of 200 ng to 1 ng of protein were resolved by SDS-PAGE using AnykD Mini-Protean TGX gels (Bio-Rad Laboratories, Hercules, CA) and transferred to nitrocellulose membranes (0.45 μm pore size; GE Healthcare, Chicago, IL). To evaluate the reactivity of each mAb, the BamA β-barrel was diluted in 8M urea in Laemmli sample buffer and incubated for 30 min at RT. A gradient of 200 ng to 1 ng of protein were resolved by SDS-PAGE using 12.5% SDS gel and transferred to nitrocellulose membranes (0.45 μm pore size). For the specific reactivity of *Pf*Trx^BamA/ECL4^ antisera against BamA ECL4, a graded amount of TbpB-LCL^BamA/ECL4^ protein (200 ng to 1 ng) was resolved by SDS-PAGE using AnykD Mini-Protean TGX gels and transferred to nitrocellulose. To assess reactivity of mouse syphilitic serum (MSS) and IRS with *Pf*Trx^BamA/ECL4^, 400 ng of the protein was immunoblotted as described above. All experimental conditions are detailed in Table S2.

### Propagation of *Tp*

The Nichols strain of *Tp* was propagated by intratesticular inoculation of adult male NZW rabbits as previously described and harvested at peak orchitis (29, 30).

### Generation of mouse syphilitic serum

Six-to eight-week-old C3H/HeJ mice were inoculated intradermally (between the scapulae), intraperitoneally, intrarectally and intragenitally (females, intravaginally; males, percutaneously in the corpus cavernosa) with 2.5×10^7^ organisms per site in 50 μl CMRL containing 20% NRS (totaling 1×10^8^ total organisms/animal). Intrarectal and intravaginal inoculations were performed with a gavage-type needle. Mice were sacrificed on day 84 post-inoculation and exsanguinated. A pool of MSS was prepared for use in opsonophagocytosis assay (detailed below).

### Macrophage preparation

Bone marrow derived macrophages (BMDM) for the murine opsonophagocytosis assay were generated as previously described (31) with the following amendments. BMDM were plated on a Millicell EZ 8-well chamber slide (Sigma-Aldrich) with 500 μl for a final concentration of 1×10^5^ cells per well and incubated overnight at 37°C. The following day, the media was replaced with fresh DMEM supplemented with 10% FBS. Rabbit peritoneal macrophages were generated using 10% protease peptone and isolated using ice-cold PBS-EDTA as previously described (30). The macrophages were plated (1×10^5^ cells/well) in 8-well BioCoat Poly-D-Lysine glass culture slide chamber slides (Corning, Corning, NY) and incubated at 37°C for 2 h. Nonadherent cells were removed by washing the monolayers twice with DMEM (30).

### Opsonophagocytosis assay

Freshly harvested *Tp* were diluted to 1×10^8^ in media alone or DMEM supplemented with 10% of normal sera, syphilitic sera, α-*Pf*Trx^BamA/ECL4^, α-*Pf*Trx^Empty^, α-Tpp17, α-TP0751 (10, 30, 31). For the murine assay, *Tp* was pre-incubated with 10 μg/ml of each mAb. After 2 hr pre-incubation at RT, *Tp* were added to macrophages at an MOI of 10:1 and incubated for 4 h at 37°C. Each stimulation condition was performed in triplicate. Following the incubation period an immunofluorescence assay (IFA) was performed and imaged to evaluate treponeme internalization as detailed below.

### Immunofluorescence for *Tp* internalization

IFA was performed as previously described (30), with the modifications in Table S2. In addition, VECTASHIELD Antifade mounting medium without DAPI (Vector Laboratories, Newark, CA) was added and samples were sealed with coverslips. Internalization of *Tp* was assessed by acquiring images of at least 100 macrophages per condition on an epifluorescence Olympus BX-41 microscope using a 40× (1.4-NA) oil immersion objective equipped with a Retiga Exi CCD camera (Q Imaging, Tucson, AZ) and the following Omega filter sets: DAPI, FITC, and rhodamine. Acquired images were processed with VisiView (version 5.0.0.7). Confocal images were acquired using Zeiss 880 and images were processed using ZEN3.5 Blue. The phagocytic index was calculated by dividing the number of internalized spirochetes by the number of cells containing internalized spirochetes and multiplying that by the number of cells containing internalized spirochetes divided by the total number of cells imaged and multiplied by 100. The percentages of macrophages containing degraded spirochetes were systematically quantified for each of the conditions studied in a blinded fashion.

### Statistical analysis

General statistical analysis was conducted using GraphPad Prism 9.5.1 (GraphPad Software, San Diego, CA). The means of the AUC from ELISA dilution curves for the *Pf*Trx^BamA/ECL4^ construct and peptides were compared to determine statistical significance by one-way ANOVA with Bonferroni’s correction for multiple comparisons. Phagocytic indexes were compared among the different stimuli. Either a paired or unpaired Student’s *t*-test (*i.e*., Mann–Whitney test or Wilcoxon test) was used for comparison across two groups. For the analysis of three or more conditions, non-parametric statistical test (Friedman’s test with a Dunnett’s multiple comparisons post-test analysis) was used for trend analysis. For each experiment, the standard error of the mean was calculated and a *p*-value < 0.05 was considered significant.

## RESULTS

### Production of mAbs targeting BamA ECL4 using Epivolve

Structural modeling of *Tp* BamA predicts that three ECLs, ECL4, ECL6, and ECL7 form a dome that covers the β-barrel’s extracellular opening (**Figure 2A**). We selected BamA ECL4 (**Figure 2A**) for production of mAbs using Epivolve (**Figure 1**) based on our previous report that it is an opsonic target in *Tp* (10) along with other studies demonstrating that Abs directed against BamA ECL4 in *E. coli* are growth limiting or bactericidal (15, 32). The ECL was divided into three overlapping peptide sequences (S1, S2 and S3), each with a centrally located, modified aspartic acid residue (**Figure 2B**). Analysis using ElliPro predicted two high scoring linear BCEs in S2 and S3 and none in S1 (**Figure 2B**). Initial rounds of panning identified phage that bound to all three modified peptides (14 hits for S1, 42 for S2 and 52 for S3). Eight phages containing a single-chain variable fragment (scFv) that bound specifically to the S2 modified peptide were advanced to the maturation step. Titration ELISAs were done to identify clones with a background-corrected signal ≥2-fold against the modified vs the native peptide (**Figure 2C**, left). To improve the binding affinity of the scFvs, scFv mutagenesis was performed using error prone PCR followed by selection against the native peptide. Hits selected for further study demonstrated a ≥ 10-and ≥ 40-fold improvement in reactivity against the modified and native peptides, respectively, compared to the parental clones and failed to bind the irrelevant peptide negative control (**Figure 2C**, right).

### Predicted CDR sequences harbor amino acid differences with the potential to impact antigen recognition

Epivolve yielded five distinct scFvs consisting of five unique heavy chains (HCs) paired with three unique light chains (LCs) (**Figure 3A**). We used abYsis (33) to determine the predicted CDR boundaries of each chain and generated separate alignments of the HC and LC sequences (**Figure 3B** and **3C**, respectively). Inspection of the alignments revealed amino acid differences in both chains potentially relevant to antigen recognition. HC1 and HC2 contain four identical substitutions in CDR2. CDR3-HC contains three substitutions: G102S in HC1, Q102H in HC2, and S107R in HC3 and HC4. LC1-CDR2 contains substitutions at positions 52 (M→R) and 55 (P→Y). CD3-LC contains A/F/S substitutions at residue 94 in all three LCs and D95K and F99M substitutions in LC1. While most of the variants were observed in the CDR regions, substitutions were identified in the framework of both chains (*e*.*g*., D10V and S13P in HC2 and LC2, respectively).

**Figure 3.**
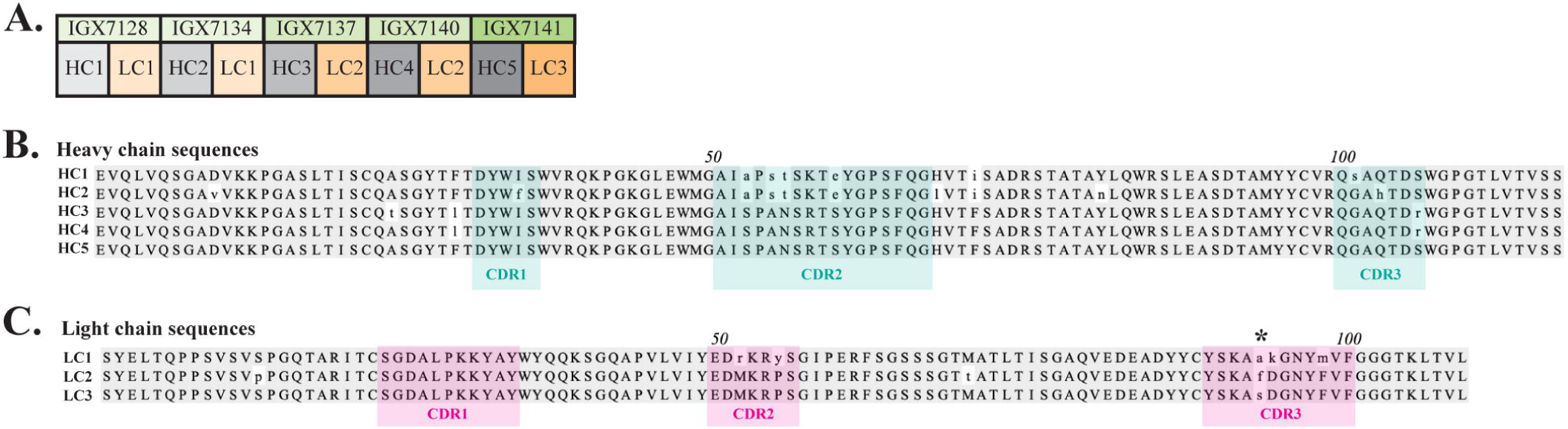
Heavy and light chains used to form full-length mAbs. (**A**) Five heavy and three light chains (HC and LC, respectively) were fused to a mouse IgG2 constant domain to form five distinct full-length mAbs. A multiple sequence alignment of the (**B**) HCs and (**C**) LCs. Grey shading represents >51% consensus identity among amino acid residues, while lower-case letters indicate mismatched residues. HC and LC CDRs predicted using abYsis (33) are denoted by cyan or pink shading, respectively. Asterisk (*) indicates a nonconserved residue in all three LC CDR3s.

### Full-length monoclonal antibodies strongly recognize BamA ECL4

The five scFvs were fused with a mouse IgG_2_ Fc backbone anticipating evaluation of their opsonic activity (see below) (34, 35). We then used ELISA to compare binding of the mAbs to the native S2 peptide and ECL4 displayed by *Pf*Trx, a scaffold protein that presents OMP ECLs in a native-like conformation (27). Each mAb demonstrated strong reactivity with both antigens (**Figure 4A**); however, based on AUC values the mAbs showed slightly greater recognition of the peptide (**Figure 4B**). By immunoblotting, four of the five mAbs detected at least 25 ng of *Pf*Trx^BamA/ECL4^, IGX7141 being the most sensitive, while the control mAb (IGX6939) failed to recognize the highest amount of protein (200 ng) (**Figure 4C**). Immunoblots against the BamA β-barrel yielded similar results (**Figure 4C**). The three mAbs with the strongest reactivity by immunoblot (IGX7137, IGX7140, and IGX7141) also showed the greatest AUC values (**Figure 4B**). Also noteworthy, the immunoblot reactivities of several mAbs compared favorably to a rat ECL4 antiserum previously demonstrated to be capable of detecting native BamA, a low abundance protein (∼200 copies/cell) (9), on the surface of intact spirochetes (**Figure 4D**) (10).

**Figure 4.**
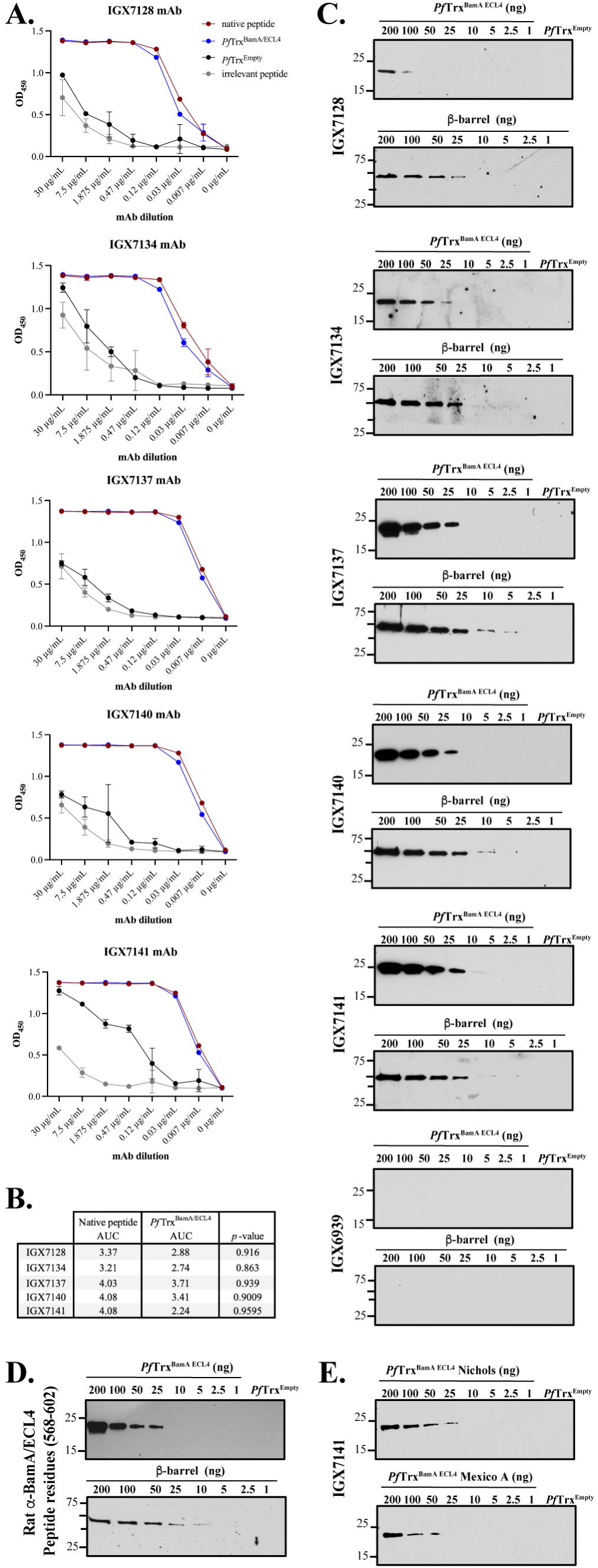
Immunoreactivity profiles of full-length of BamA ECL4 mAb. (**A**) Titration ELISAs of the mAbs against native and irrelevant peptides (magenta and grey, respectively) and *Pf*Trx^BamA/ECL4^ and *Pf*Trx ^Empty^ (blue and black, respectively). (**B**) mAb AUC values calculated for native peptide and *Pf*Trx^BamA/ECL4^. (**C**) Immunoblot reactivities against graded nanogram amounts of *Pf*Trx ^BamA/ECL4^ or the BamA β-barrel. *Pf*Trx^Empty^ (200 ng) and nonspecific mAb (IGX6939) served as specificity controls. Immunoblots were done using mAbs at 4 μg/ml. (**D**) A rat polyclonal BamA ECL4 (residues 568-602) antiserum (diluted 1:1000) was used as a comparator (10). (**E**) Immunoblot reactivity of IGX7141 mAb (4 μg/ml) against graded nanogram amounts of Nichols (top) and Mexico A *Pf*Trx^BamA/ECL4^ (bottom) variants.

We previously reported that ECL4 harbors an immunodominant epitope in which substitution of glutamine in the Mexico A strain for leucine at position 594 in the Nichols strain markedly diminishes Ab recognition (10, 29). This residue is the last amino acid in the S2 peptide (**Figure 2B**). We next sought to determine if this substitution impacts recognition of ECL4 by the mAbs. IGX7141, the strongest reactor, exhibited slightly diminished recognition of the Mexico A ECL4 variant displayed by the *Pf*Trx scaffold (**Figure 4E**).

### Identification of an opsonic BamA ECL4 monoclonal Ab

Macrophage-mediated opsonophagocytosis is considered to be critical for treponemal clearance (36), and *ex vivo* opsonic activity is widely considered to be a correlate of protective immunity (5, 37). Opsonophagocytosis assays with *Tp* are typically done with rabbit sera and rabbit peritoneal macrophages (30, 37); however, we previously demonstrated that opsonic activity also can be assessed using sera from mice infected with *Tp* (MSS) and murine bone marrow-derived macrophages (31). We employed the latter assay to evaluate the opsonic activity of the five mAbs. As a positive control we collected and pooled sera from five mice infected with *Tp* for 84 days, a time point known to elicit strongly opsonic Abs (31). As a presumptive positive control, we pooled mouse polyclonal antisera generated against *Pf*Trx^BamA/ECL4^ and confirmed the presence of ECL-specific Abs by immunoblotting against a heterologous scaffold, TbpB-LCL (*Neisseria meningitidis* <u>T</u>ransferrin-<u>B</u>inding <u>P</u>rotein <u>B</u>, ‘loopless’ <u>C</u>-lobe) (28), displaying ECL4 (**Figure 5A**). The polyclonal ECL4 antisera recognized the S2 peptide as well as peptides S1 and S3 (**Figure 5B**). In addition to normal mouse serum (NMS) and mouse *Pf*Trx^Empty^ antisera, as negative controls we generated murine antisera against the periplasmic lipoproteins Tpp17 and TP0751 (**Figure S3**) (7, 30). Previously, spirochete uptake has been calculated as a percentage of macrophages containing *Tp* (30, 38). Our use of confocal microscopy enabled us to devise an improved ‘phagocytic index’ that quantifies both the number of macrophages with ingested organisms and the number of treponemes phagocytosed per cell.

**Figure 5.**
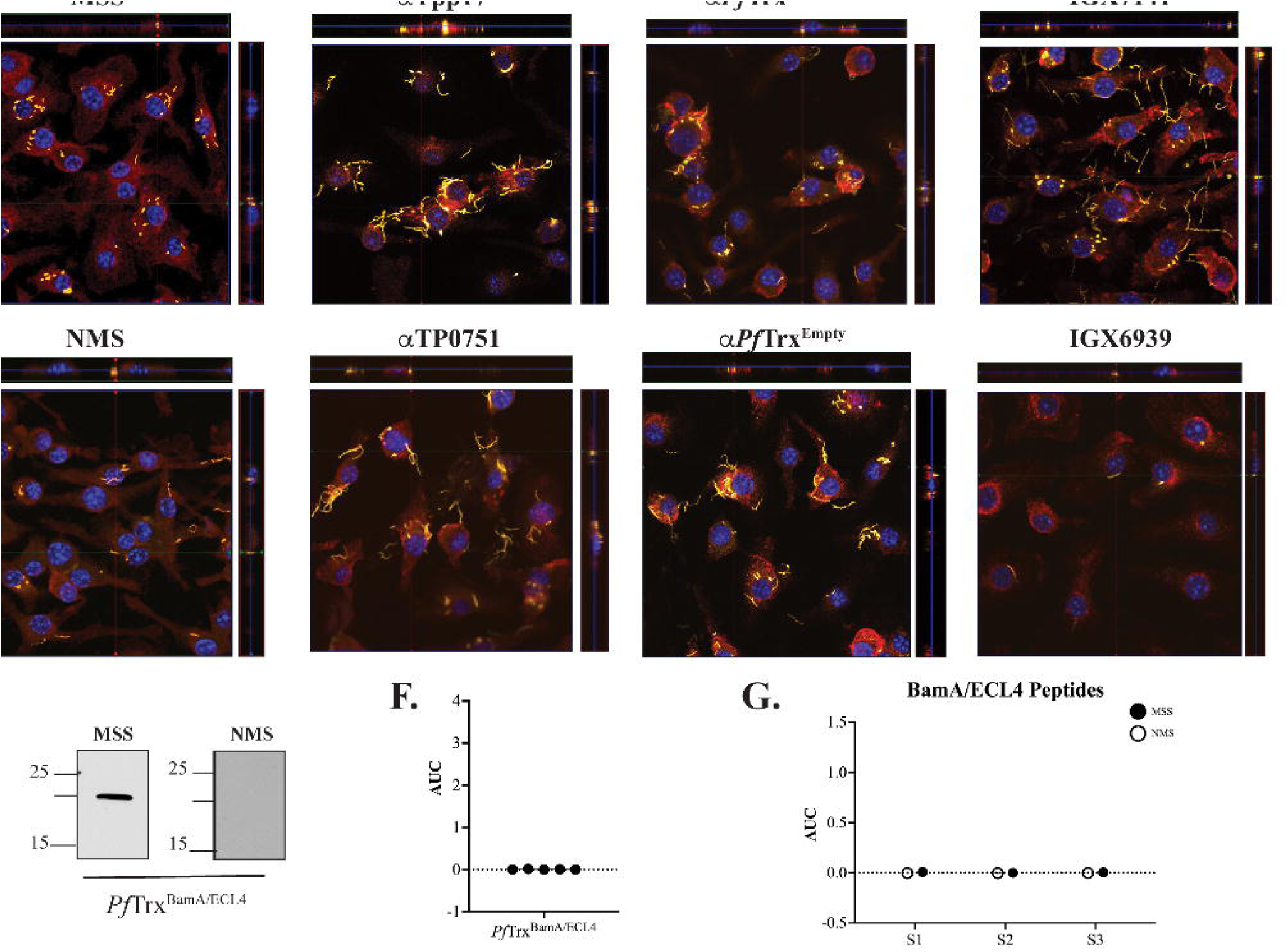
Identification of an opsonic BamA ECL4 mAb. (**A**) Immunoblot reactivities of pooled sera (diluted 1:1,000) from five mice hyperimmunized with *Pf*Trx^BamA/ECL4^ against graded nanogram amounts of TbpB-LCL^BamA/ECL4^. (**B**) ELISA reactivity of murine *Pf*Trx^BamA/ECL4^ antisera or NMS with native S1, S2, and S3 peptides represented as AUC values. (**C**) Freshly extracted *Tp* were pre-incubated with 10% heat-inactivated NMS, pooled MSS, mouse antisera to *Pf*Trx^BamA/ECL4^, *Pf*Trx^Empty^, TP0751 or Tpp17 or 10 μg/ml of the individual mAbs followed by incubation with murine BMDMs for 4 h at an MOI 10:1. Phagocytic indices were determined as described in Materials and Methods. Asterisks show significant differences with *p*-values of ≤0.05, ≤0.01 or <0.0001. (**D**) Each representative confocal micrograph is a composite of 9-12 consecutive Z-stack planes with labeling of *Tp*, plasma membranes and nuclei shown in green, red and blue, respectively. (**E**) Immunoblot reactivity of pooled MSS and NMS (diluted 1:250) against 200 ng of *Pf*Trx^BamA/ECL4^. ELISA reactivity (AUC values) of pooled MSS against (**F**) *Pf*Trx^BamA/ECL4^ and (**G**) the S1, S2 and S3 peptides.

As shown in **Figure 5C** and reported previously (31), MSS exhibited robust opsonic activity, whereas spirochetes pre-incubated with NMS bound to the surface of macrophages but were poorly internalized. Also consistent with a previous report using the rabbit assay (30), mouse Abs directed against Tpp17 and TP0751 exhibited background levels of phagocytosis (**Figure 5C** and **5D**). In contrast, spirochetes pre-incubated with the pooled ECL4 antisera were internalized at levels well above background (*p* = 0.031). Of the five mAbs, only mAb IGX7141 displayed significant opsonic activity (*p* = 0.003); its opsonic activity was comparable to that of the ECL4 antisera (**Figure 5C** and **5D**). Interestingly, the MSS showed a markedly different reactivity profile for ECL4 than either the mAbs or the polyclonal antisera; it reacted poorly by immunoblot (**Figure 5E**) and failed to recognize *Pf*Trx^BamA/ECL4^ and all three peptides by ELISA (**Figure 5F** and **5G**).

### Immune rabbit serum lacks antibodies to the subdominant BamA ECL4 epitope

As noted above, the rabbit opsonophagocytosis assay is the conventional method for assessing opsonic activity for *Tp*. We, therefore, next sought to determine how the opsonization and antigenicity data obtained in the murine assay correlated with results obtained with the rabbit system. A rabbit antiserum generated using *Pf*Trx^BamA/ECL4^ displayed similar immunoblot reactivity to TbpB-LCL^BamA/ECL4^ as its mouse counterpart (**Figure 6A**). Notably, compared to the mouse ECL4 antisera (**Figure 5B**), the AUC values of the rabbit antiserum for all three peptides were substantially greater (**Figure 6B**). As in the mouse assay, as additional negative controls, we included previously characterized rabbit antisera against Tpp17 and TP0751 (30). As shown in **Figure 6C** and **6D**, sera from five immune rabbits and the polyclonal ECL4 antiserum showed strong opsonic activity. The greater peptide ELISA values of the rabbit ECL4 antiserum vs. the mouse likely explains its greater opsonic activity (compare **Figure 5C** and **5D** with **Figure 6C** and **6D**). Four of the five immune sera reacted strongly with *Pf*Trx^BamA/ECL4^ by immunoblot (**Figure 6E**); one of these (IRS 114) failed to recognize ECL4 by ELISA (**Figure 6F**). Three of the four immunoblot positive immune sera were nonreactive with all three peptides, whereas one (IRS 112) recognized peptides S2 and S3 (**Figure 6G**), albeit more poorly than the rabbit ECL4 antiserum. IRS 113 was nonreactive in all three assays.

**Figure 6.**
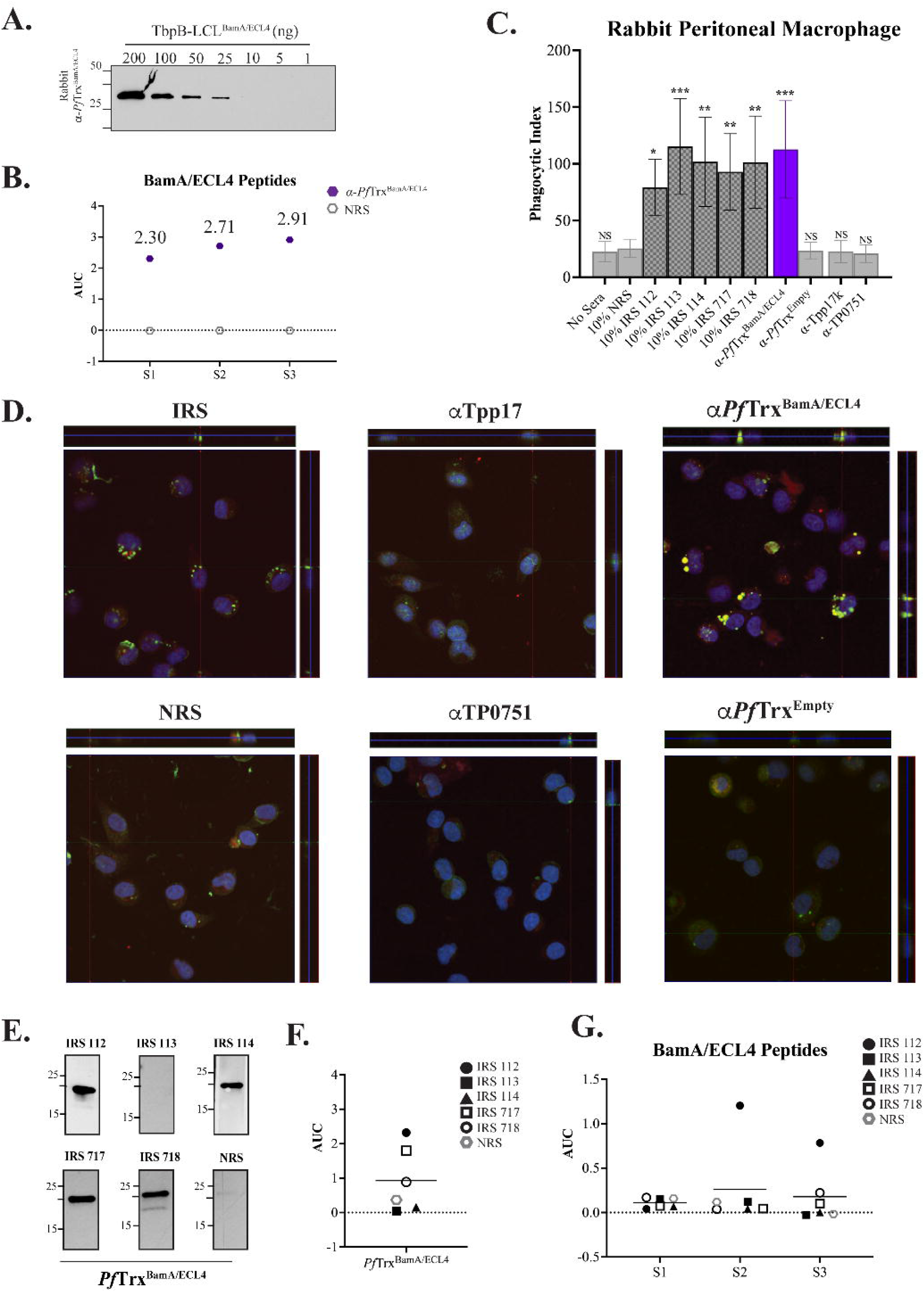
Absence of antibodies to the subdominant BamA ECL4 epitope in immune rabbit serum. (**A**) Immunoblot reactivities of sera (diluted 1:1,000) from rabbits hyperimmunized with *Pf*Trx^BamA/ECL4^ against graded nanogram amounts of TbpB-LCL^BamA/ECL4^. (**B**) ELISA reactivity of rabbit *Pf*Trx^BamA/ECL4^ antisera or NMS with native S1, S2, and S3 peptides represented as AUC values. (**C**) Freshly extracted *Tp* were pre-incubated with 10% heat-inactivated NRS, five individual IRS, or sera from rabbits hyperimmunized with *Pf*Trx^BamA/ECL4^, *Pf*Trx^Empty^, TP0751 or Tpp17 followed by incubation with rabbit peritoneal macrophages for 4 h at an MOI 10:1. Phagocytic indices were determined as described in Materials and Methods. Asterisks show significant differences with *p*-values of ≤0.05, ≤0.01, or ≤0.001. (**D**) Each representative confocal micrograph is a composite of 9-12 consecutive Z-stack planes with labeling of *Tp*, plasma membranes and nuclei shown in green, red and blue, respectively. (**E**) Immunoblot reactivity of individual IRS and NRS (diluted 1:250) against 200 ng of *Pf*Trx^BamA/ECL4^. ELISA reactivity (AUC values) of IRS against (**F**) *Pf*Trx^BamA/ECL4^ and (**G**) the S1, S2 and S3 peptides.

## DISCUSSION

The current conception of protective immunity is that spirochete clearance is driven by opsonophagocytosis and that production of so-called “functional” Abs must be paired with cellular responses to activate professional phagocytes, particularly macrophages (38, 39, 40, 41). The primary targets for opsonic Abs in syphilitic sera are presumed to be the ECLs of rare OMPs. Based on recently refined 3D structural models for *Tp* OMPs, it is now possible to generate opsonic polyclonal and mAbs directed against selected ECLs. Whereas polyclonal Abs will identify ECLs capable of serving as opsonic targets, mAbs will precisely define the paratope-epitope interactions required for opsonic activity. Herein, we generated a mAb that recognizes an opsonic epitope on ECL4 of *Tp* BamA. Abs against this epitope are not strongly elicited during natural infection; on the other hand, hyperimmunization with ECL4 stimulates the immune system to outflank this “antigenic barrier” giving rise to Abs that detect what appears to be a subdominant epitope.

Epivolve technology yielded five mAbs that recognized a centrally located (S2) BamA ECL4 peptide. All five mAbs reacted by ELISA with both the native peptide and the ECL presented in a “native-like” conformation within the context of a *Pf*Trx scaffold. Immunoblot reactivity with *Pf*Trx^BamA/ECL4^ and the β-barrel confirmed that the mAbs recognize a linear epitope. Taken together, these data demonstrate that a linear epitope can be displayed in an Ab accessible manner when a large, conformationally dynamic polypeptide (in this case 35 residues) is constrained at both ends as occurs within the native ECL. While the reactivities of the five mAbs were similar, they were not identical; three of the five mAbs (IGX7137, IXG7140 and IGX7141) demonstrated higher affinity. These differences in reactivity likely can be explained by the amino acid variances found in CDR2 of HC1 and HC2 and the residue differences observed in CDR2 and CDR3 of LC1. Interestingly, two of three strongly reactive mAbs (IGX7137 and IXG7140) share LC2. While LC3 is highly similar to LC2, the single amino acid difference observed at position 94 of CDR3 presumably is responsible for the unique opsonic activity of IGX7141. Importantly, each of the IgGs have different heavy chains and could also significantly contributes to Ab binding and may be responsible for the reactivity and/or opsonic activity.

To evaluate functional activities of the mAbs, we utilized our previously described opsonophagocytosis assay employing murine macrophages. In the current study, we used C3H/HeJ mice, rather than the C57BL/6 strain, based on prior publications (42) and our recent experience that the former produce higher Ab titers following hyerimmunization with *Pf*Trx-scaffolded ECLs. At the outset, we confirmed that *Tp* infection of C3H/HeJ mice elicits strongly opsonic Abs. It is worth noting that spirochetes pre-incubated with NMS readily bound to the surface of macrophages but were not internalized. The lack of opsonic activity observed with mouse Abs to Tpp17 and TP0751 confirmed previous results obtained with rabbit antisera (30) and is in accord with studies localizing both lipoproteins to the periplasmic space (30, 43). As noted above, the opsonic activity of IGX7141 appears to be attributable to the single amino acid substitution in CDR3 of the LC. Studies have shown that CDR3 is a critical domain for antigen recognition (44). Previously noted differences in the frameworks of the mAbs also may impact how they engage their cognate epitope. A more detailed study of binding perimeters is needed to better understand the affinities of these five mAbs. It is noteworthy that the opsonic activity of mAb IGX7141 was equivalent to the mouse polyclonal antisera. Expansion of the study to include the conventional rabbit opsonophagocytosis assay allowed, for the first-time, a comparison of mouse and rabbit ECL-specific antisera. Unlike the mouse ECL4 antisera, the rabbit antiserum showed a comparable level of internalization to IRS. This difference appears to be attributable to the stronger reactivity of the rabbit antiserum with the ECL peptides rather than any fundamental difference between the two assays. Regardless, both assays underscore that hyperimmunization against a single ECL can elicit strong opsonic activity.

A striking observation is that infection with *Tp* often does not elicit opsonic Abs against the target recognized by the ECL4 mAbs. Thus, while immune serum harbors Abs against BamA ECL4, their contribution to the overall opsonic activity of syphilitic serum remains unclear. More broadly, these findings raise the possibility that *Tp* diverts the host immune response away from subdominant opsonic ECL epitopes as part of its strategy for stealth pathogenicity (45). Epivolve is divorced from the natural immune response and allows for the generation of mAbs which can target subdominant epitopes. Hyperimmunization with an ECL displayed on *Pf*Trx also was able to overcome this immunologic barrier, although it remains to be determined if this will be the case for other ECLs (46). Importantly, the opsonic mAb IGX7141 recognized both ECL4 variants, supporting that Abs directed against this subdominant epitope can sidestep the *Tp*’s attempt at immune evasion through antigenic variation of a surface exposed epitope on circulating strains. Here we demonstrate the utility of mAbs generated outside the immune system to identify potentially protective ECL epitopes that would be missed by relying solely on screening approaches based upon the natural Ab response to *Tp*.

## Author Contributions

MF, KD, SM, MC, MM, JR, MW and KH contributed to conception and design of the study. MF, KD and KH organized the database. KD and KH performed the statistical analysis. JR and KH wrote the first draft of the manuscript. MF, KD, and MC wrote sections of the manuscript. All authors contributed to manuscript revision, read, and approved the submitted version.

## Acknowledgments

We thank Ms. Morgan LeDoyt, Ms. Crystal Vicente and Mr. Kemar Edwards (UConn Health, USA) for their expert technical support.

## FIGURE LEGENDS

### SUPPLEMENTARY FIGURE LEGENDS

**Figure S1. Library vector template.** The parental vector was modified to contain four *Eco*29kI restriction endonuclease sites and amber (5’-TAG-3’) stop codons within the CDRs targeted for mutagenesis. For library generation, amino acid stretches varying from six to 22 residues were incorporated into LC and HC CDR3s. Colored bars represent designate primer binding sites.

**Figure S2. Production of high-titer, fully recombinant Ab libraries.** (**A**) *In vivo* restriction using *E. coli* AXE688 [TG1 (*eco*29KI.RM)] to produce recombinant libraries (21, 22). Parental plasmids carrying *Eco*29kI sites within the complementarity determining regions (CDRs) of the scFv are cleaved by *Eco*29kI expressed in the AXE688 cells. (**B**) *In vivo* selection using saturating DNA. Super-saturating concentrations of plasmid DNA were used generate large recombinant libraries. Competent cells take up multiple plasmids under DNA saturating conditions using AXE688, thereby resulting in transformed cells with a higher proportion of totally recombinant clones.

**Figure S3. Reactivity of mouse Tpp17 and TP0751 antisera.** Reactivity of sera (diluted 1:1,000) from mice hyperimmunized with Tpp17 or TP0751 by immunoblot analysis against graded nanogram amounts of (**A**) Tpp17 or (**B**) TP0751.

## SUPPLEMENTARY TABLES

**Table S1. Primers**

**Table S2. Immunological assay conditions**

## REFERENCES

1. Hook EWR. Syphilis. Lancet. 2017;389(10078):1550–7.

2. Radolf JD, Tramont EC, Salazar JC. Syphilis (Treponema pallidum). In: Mandell GL, Dolin R, Blaser MJ, editors. Mandell, Douglas and Bennett’s Principles and Practice of Infectious Diseases. 9 ed. Philadelphia: Churchill Livingtone Elsevier; 2019. p. 2865–92.

3. Patton ME, Su JR, Nelson R, Weinstock H, Centers for Disease C, Prevention. Primary and secondary syphilis--United States, 2005-2013. MMWR Morb Mortal Wkly Rep. 2014;63(18):402-6.

4. Peeling RW, Mabey D, Kamb ML, Chen XS, Radolf JD, Benzaken AS. Syphilis. Nat Rev Dis Primers. 2017;3:17073.

5. Radolf JD, Lukehart SA. Immunology of Syphilis. In: Radolf JD, Lukehart SA, editors. Pathogenic Treponemes: Cellular and Molecular Biology. Norfolk, UK: Caister Academic Press; 2006. p. 285–322.

6. Hawley KL, Montezuma-Rusca JM, Delgado KN, Singh N, Uversky VN, Caimano MJ, et al. Structural modeling of the *Treponema pallidum* outer membrane protein repertoire: a road map for deconvolution of syphilis pathogenesis and development of a syphilis vaccine. J Bacteriol. 2021;203(15):e0008221.

7. Cox DL, Luthra A, Dunham-Ems S, Desrosiers DC, Salazar JC, Caimano MJ, et al. Surface immunolabeling and consensus computational framework to identify candidate rare outer membrane proteins of *Treponema pallidum*. Infect Immun. 2010;78:5178–94.

8. Cameron CE, Lukehart SA, Castro C, Molini B, Godornes C, Van Voorhis WC. Opsonic potential, protective capacity, and sequence conservation of the *Treponema pallidum* subspecies *pallidum* Tp92. J Infect Dis. 2000;181(4):1401–13.

9. Desrosiers DC, Anand A, Luthra A, Dunham-Ems SM, LeDoyt M, Cummings MA, et al. TP0326, a *Treponema pallidum* ®-barrel assembly machinery A (BamA) orthologue and rare outer membrane protein. Mol Microbiol. 2011;80(6):1496–515.

10. Luthra A, Anand A, Hawley KL, LeDoyt M, La Vake CJ, Caimano MJ, et al. A homology model reveals novel structural features and an immunodominant surface loop/opsonic target in the *Treponema pallidum* BamA ortholog TP_0326. J Bacteriol. 2015;197(11):1906–20.

11. Andreano E, Seubert A, Rappuoli R. Human monoclonal antibodies for discovery, therapy, and vaccine acceleration. Curr Opin Immunol. 2019;59:130–4.

12. Taylor PC, Adams AC, Hufford MM, de la Torre I, Winthrop K, Gottlieb RL. Neutralizing monoclonal antibodies for treatment of COVID-19. Nat Rev Immunol. 2021;21(6):382–93.

13. Mazur NI, Terstappen J, Baral R, Bardaji A, Beutels P, Buchholz UJ, et al. Respiratory syncytial virus prevention within reach: the vaccine and monoclonal antibody landscape. Lancet Infect Dis. 2022.

14. Cai F, Chen WH, Wu W, Jones JA, Choe M, Gohain N, et al. Structural and genetic convergence of HIV-1 neutralizing antibodies in vaccinated non-human primates. PLoS Pathog. 2021;17(6):e1009624.

15. Wang H, Chen D, Lu H. Anti-bacterial monoclonal antibodies: next generation therapy against superbugs. Appl Microbiol Biotechnol. 2022;106(11):3957–72.

16. Jones SA, Marchitto KS, Miller JN, Norgard MV. Monoclonal antibody with hemagglutination, immobilization, and neutralization activities defines an immunodominant, 47,000 mol wt, surface-exposed immunogen of *Treponema pallidum* (Nichols). J Exp Med. 1984;160(5):1404–20.

17. Robertson SM, Kettman JR, Miller JN, Norgard MV. Murine monoclonal antibodies specific for virulent *Treponema pallidum* (Nichols). Infect Immun. 1982;36(3):1076–85.

18. Saunders JM, Folds JD. Development of monoclonal antibodies that recognize *Treponema pallidum*. Infect Immun. 1983;41(2):844–7.

19. Cox DL, Chang P, McDowall AW, Radolf JD. The outer membrane, not a coat of host proteins, limits antigenicity of virulent *Treponema pallidum*. Infect Immun. 1992;60(3):1076–83.

20. Cox DL, Akins DR, Porcella SF, Norgard MV, Radolf JD. *Treponema pallidum* in gel microdroplets: a novel strategy for investigation of treponemal molecular architecture. Mol Microbiol. 1995;15(6):1151–64.

21. Holland EG, Acca FE, Belanger KM, Bylo ME, Kay BK, Weiner MP, et al. In vivo elimination of parental clones in general and site-directed mutagenesis. J Immunol Methods. 2015;417:67–75.

22. Holland EG, Buhr DL, Acca FE, Alderman D, Bovat K, Busygina V, et al. AXM mutagenesis: an efficient means for the production of libraries for directed evolution of proteins. J Immunol Methods. 2013;394(1-2):55–61.

23. Zakharova MV, Beletskaya IV, Kravetz AN, Pertzev AV, Mayorov SG, Shlyapnikov MG, et al. Cloning and sequence analysis of the plasmid-borne genes encoding the Eco29kI restriction and modification enzymes. Gene. 1998;208(2):177–82.

24. Dan Z, Tan Z, Xia H, Wu G. Construction and expression of D-dimer and GPIIb/IIIa single-chain bispecific antibody. Exp Ther Med. 2013;6(2):552–6.

25. Zhao Q, Buhr D, Gunter C, Frenette J, Ferguson M, Sanford E, et al. Rational library design by functional CDR resampling. N Biotechnol. 2018;45:89–97.

26. Ponomarenko J, Bui HH, Li W, Fusseder N, Bourne PE, Sette A, et al. ElliPro: a new structure-based tool for the prediction of antibody epitopes. BMC Bioinformatics. 2008;9:514.

27. Delgado KN, Montezuma-Rusca JM, Orbe IC, Caimano MJ, La Vake CJ, Luthra A, et al. Extracellular loops of the *Treponema pallidum* FadL orthologs TP0856 and TP0858 elicit IgG antibodies and IgG(+)-specific B-cells in the rabbit model of experimental syphilis. mBio. 2022;13(4):e0163922.

28. Fegan JE, Calmettes C, Islam EA, Ahn SK, Chaudhuri S, Yu RH, et al. Utility of hybrid transferrin binding protein antigens for protection against pathogenic *Neisseria* species. Front Immunol. 2019;10:247.

29. Kumar S, Caimano MJ, Anand A, Dey A, Hawley KL, LeDoyt ME, et al. Sequence variation of rare outer membrane protein beta-barrel domains in clinical strains provides insights into the evolution of *Treponema pallidum* subsp. *pallidum*, the syphilis spirochete. mBio. 2018;9(3).

30. Luthra A, Montezuma-Rusca JM, La Vake CJ, LeDoyt M, Delgado KN, Davenport TC, et al. Evidence that immunization with TP0751, a bipartite *Treponema pallidum* lipoprotein with an intrinsically disordered region and lipocalin fold, fails to protect in the rabbit model of experimental syphilis. PLoS Pathog. 2020;16(9):e1008871.

31. Silver AC, Dunne DW, Zeiss CJ, Bockenstedt LK, Radolf JD, Salazar JC, et al. MyD88 deficiency markedly worsens tissue inflammation and bacterial clearance in mice infected with *Treponema pallidum*, the agent of syphilis. PLoS One. 2013;8(8):e71388.

32. Vij R, Lin Z, Chiang N, Vernes JM, Storek KM, Park S, et al. A targeted boost-and-sort immunization strategy using *Escherichia coli* BamA identifies rare growth inhibitory antibodies. Sci Rep. 2018;8(1):7136.

33. Swindells MB, Porter CT, Couch M, Hurst J, Abhinandan KR, Nielsen JH, et al. abYsis: integrated antibody sequence and structure-management, analysis, and prediction. J Mol Biol. 2017;429(3):356–64.

34. Dekkers G, Bentlage AEH, Stegmann TC, Howie HL, Lissenberg-Thunnissen S, Zimring J, et al. Affinity of human IgG subclasses to mouse Fc gamma receptors. MAbs. 2017;9(5):767–73.

35. Collins AM. IgG subclass co-expression brings harmony to the quartet model of murine IgG function. Immunol Cell Biol. 2016;94(10):949–54.

36. Lafond RE, Lukehart SA. Biological basis for syphilis. Clin Microbiol Rev. 2006;19(1):29–49.

37. Lukehart SA. Scientific monogamy: thirty years dancing with the same bug: 2007 Thomas Parran Award Lecture. Sex Transm Dis. 2008;35(1):2–7.

38. Hawley KL, Cruz AR, Benjamin SJ, La Vake CJ, Cervantes JL, LeDoyt M, et al. IFN© enhances CD64-potentiated phagocytosis of *Treponema pallidum* opsonized with human syphilitic serum by human macrophages. Front Immunol. 2017;8:1227.

39. Cruz AR, Ramirez LG, Zuluaga AV, Pillay A, Abreu C, Valencia CA, et al. Immune evasion and recognition of the syphilis spirochete in blood and skin of secondary syphilis patients: two immunologically distinct compartments. PLoS Negl Trop Dis. 2012;6(7):e1717.

40. Marra CM, Tantalo LC, Sahi SK, Dunaway SB, Lukehart SA. Reduced *Treponema pallidum*-specific opsonic antibody activity in HIV-infected patients with syphilis. J Infect Dis. 2016;213(8):1348–54.

41. Batonick M, Holland EG, Busygina V, Alderman D, Kay BK, Weiner MP, et al. Platform for high-throughput antibody selection using synthetically-designed antibody libraries. N Biotechnol. 2016;33(5 Pt A):565–73.

42. Gwinn WM, Johnson BT, Kirwan SM, Sobel AE, Abraham SN, Gunn MD, et al. A comparison of non-toxin vaccine adjuvants for their ability to enhance the immunogenicity of nasally-administered anthrax recombinant protective antigen. Vaccine. 2013;31(11):1480–9.

43. Radolf JD, Kumar S. The *Treponema pallidum* outer membrane. Curr Top Microbiol Immunol. 2018;415:1–38.

44. Rock EP, Sibbald PR, Davis MM, Chien YH. CDR3 length in antigen-specific immune receptors. J Exp Med. 1994;179(1):323–8.

45. Radolf JD, Deka RK, Anand A, Smajs D, Norgard MV, Yang XF. *Treponema pallidum*, the syphilis spirochete: making a living as a stealth pathogen. Nat Rev Microbiol. 2016.

46. Lindahl G. Subdominance in antibody responses: implications for vaccine development. Microbiol Mol Biol Rev. 2020;85(1).

