## Supplementary figures and images for "Use of Epivolve phage display to generate a monoclonal antibody with opsonic activity directed against a subdominant epitope on extracellular loop 4 of *Treponema pallidum* BamA (TP0326)"

### Supplementary Figure 1

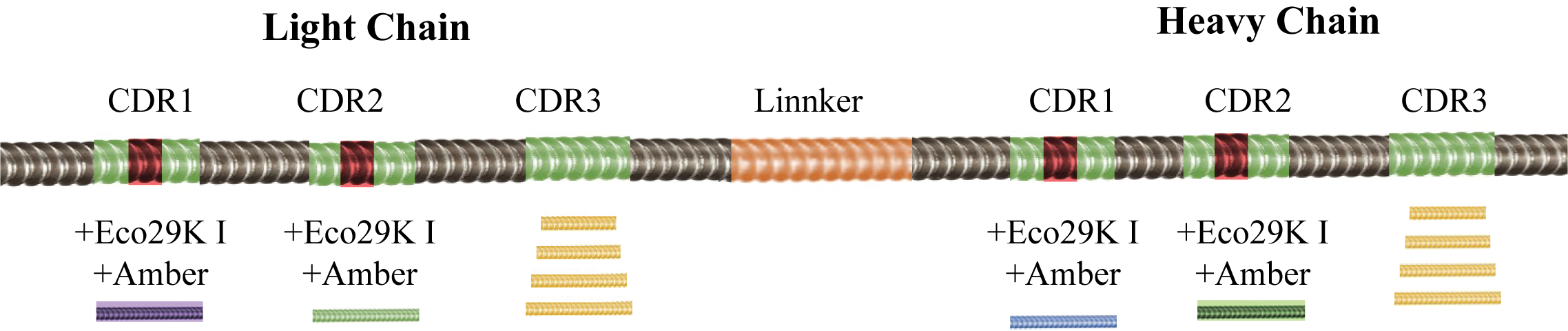

### Supplementary Figure 2

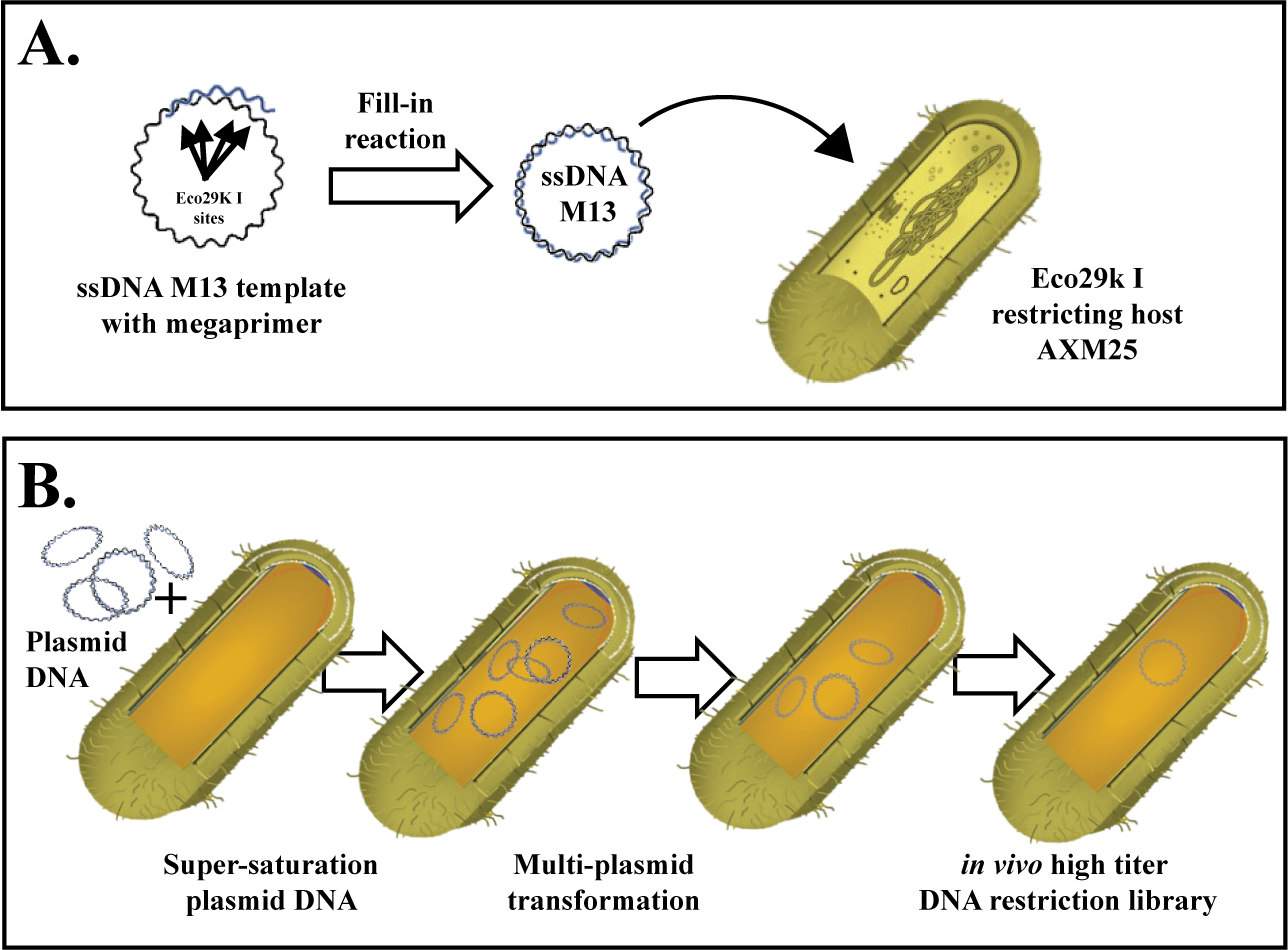

### Supplementary Figure 3

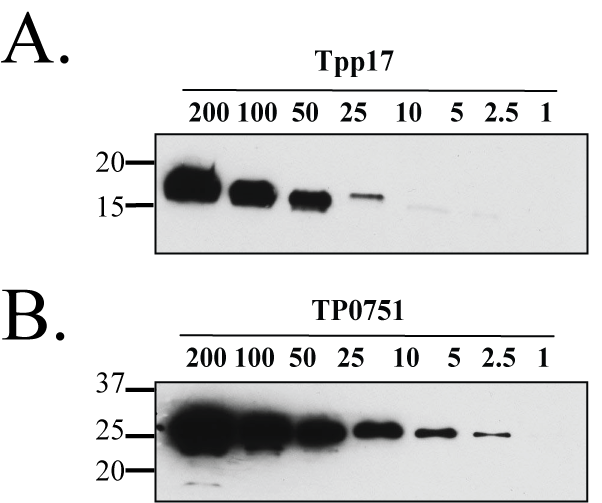
