## Supplementary Table 1 for "Use of Epivolve phage display to generate a monoclonal antibody with opsonic activity directed against a subdominant epitope on extracellular loop 4 of *Treponema pallidum* BamA (TP0326)"

| Protein | Name | Primer name | Description | Oligo Sequence (5' to 3') |
|---|---|---|---|---|
| **BamA** | *Pf*Trx$^{BamA/ECL4}$, (Nichols) | *Pf*Trx$^{BamA/ECL4}$ –FW | Amplification of BamA ECL4 $^{568\text{-}602}$, Nichols | CGCGCGGCAGCCATATGAGCAGCGGCATTATCGAG |
| | | *Pf*Trx$^{BamA/ECL4}$ –RV | | GGTGGTGGTGCTCGAGTTACTGTGCCACTCGATC |
| | *Pf*Trx$^{BamA/ECL4}$, (Mexico A) | *Pf*Trx$^{BamA/ECL4}$ MexA–FW | Amplification of BamA ECL4 $^{568\text{-}602}$, Mexico A | GCCGTTCGATCAAACCGTGAAAG |
| | | *Pf*Trx$^{BamA/ECL4}$ MexA–RV | | TGGTTGTTGTCCTTGTCG |
| **TbpB loopless C-lobe** | TbpB-LCL$^{BamA/ECL4}$ | TbpB28nt-F | Amplification of BamA ECL4 $^{568\text{-}602}$, Nichols | CAGCCATATGGCTAGCGGCAGCAGCAGCGAAAACA |
| | | TbpB-ntR | | GGTGGTGGTGCTCGAGTCATTGCACCGGCTGCTGACG |
