## Supplementary Table 2 for "Use of Epivolve phage display to generate a monoclonal antibody with opsonic activity directed against a subdominant epitope on extracellular loop 4 of *Treponema pallidum* BamA (TP0326)"

| Assay | Species | Antigen | First Blocking | Primary Ab | Second Blocking | Secondary Ab | Cell Markers |
|---|---|---|---|---|---|---|---|
| ELISA | Mouse | BamA ECL4 peptide | PBS containing 15% goat serum, 0.005% Tween 20 and 0.05% sodium azide; 1 h at RT | IGX mAbs (30 µg/ml); 4-fold serial dilution in 1% BSA, 1.5 h at RT | | HRP conjugated goat α-mouse Ig (1:10,000); 1 h at RT | |
| | | *Pf*Trx$^{BamA/ECL4}$ | PBS containing 15% goat serum, 0.005% Tween 20 and 0.05% sodium azide; 1 h at RT | IGX mAbs (30 µg/ml); 4-fold serial dilution in 5% BSA, 1 h at RT | | HRP conjugated goat α-mouse Ig (1:10,000); 1 h at RT | |
| | | BamA ECL4 peptide | PBS containing 15% goat serum, 0.005% Tween 20 and 0.05% sodium azide; 1 h at RT | α-*Pf*Trx$^{BamA/ECL4}$ (1:20); 2-fold serial dilution in 5% BSA, 1.5 h at RT | | HRP conjugated goat α-mouse Ig (1:10,000); 1 h at RT | |
| | Rabbit | *Pf*Trx$^{BamA/ECL4}$ | PBS containing 15% goat serum, 0.005% Tween 20 and 0.05% sodium azide; 1 h at RT | IRS (1:20); 2-fold serial dilution in 1% BSA, 1 h at RT | | HRP conjugated goat α-rabbit Ig (1:10,000); 1 h at RT | |
| | | BamA ECL4 peptide | PBS containing 15% goat serum, 0.005% Tween 20 and 0.05% sodium azide; 1 h at RT | α-*Pf*Trx$^{BamA/ECL4}$ (1:20); 2-fold serial dilution in 5% BSA, 1.5 h at RT | | HRP conjugated goat α-rabbit Ig (1:10,000); 1 h at RT | |
| Immunoblot | Mouse | *Pf*Trx$^{BamA/ECL4}$ | PBS containing 5% nonfat dry milk and 0.1% Tween 20; 1 h at RT | mAbs (4 µg/ml), ON at 4°C | | HRP-conjugated goat α-mouse Ig (1:10,000); 1 h at RT | |
| | | TbpB-LCL$^{BamA/ECL4}$ | PBS containing 5% nonfat dry milk and 0.1% Tween 20; 1 h at RT | α-*Pf*Trx$^{BamA/ECL4}$ (1:1000), ON at 4°C | | HRP-conjugated goat α-mouse Ig (1:30,000); 1 h at RT | |
| | | *Pf*Trx$^{BamA/ECL4}$ | PBS containing 5% nonfat dry milk and 0.1% Tween 20; 1 h at RT | MSS and NMS (1:250), ON at 4°C | | HRP-conjugated goat α-mouse Ig (1:30,000); 1 h at RT | |
| | Rat | *Pf*Trx$^{BamA/ECL4}$ | PBS containing 5% nonfat dry milk and 0.1% Tween 20; 1 h at RT | α-BamA/ECL4 (1:500), ON at 4°C | | HRP-conjugated goat α-rat Ig (1:30,000); 1 h at RT | |
| | Rabbit | TbpB-LCL$^{BamA/ECL4}$ | PBS containing 5% nonfat dry milk and 0.1% Tween 20; 1 h at RT | α-*Pf*Trx$^{BamA/ECL4}$ (1:1000), ON at 4°C | | HRP-conjugated goat α-rabbit Ig (1:30,000); 1 h at RT | |
| | | *Pf*Trx$^{BamA/ECL4}$ | PBS containing 5% nonfat dry milk and 0.1% Tween 20; 1 h at RT | IRS and NRS (1:250), ON at 4°C | | HRP-conjugated goat α-rabbit Ig (1:30,000); 1 h at RT | |
| IFA | Mouse | *Tp* and macrophages | 5% BSA in PBS; 1 h at RT | Rabbit anti-*Tp* (1:100), ON at 4°C | | α-rabbit IgG TexasRed (1:500); 1 h at RT | Phalloidin AF488 (1:10); 30 min Cholera Toxin AF647 (1:500); 30 min DAPI (1:1000); 10 min; All at RT |
| | Rabbit | | CMRL 10% NGS; 1 h at RT | MSS (1:25), ON at 4°C | CMRL 10% NGS; 1 h at RT | α-mouse IgG AF488 (1:500); 1 h at RT | Cholera Toxin AF647 (1:500); 30 min DAPI (1:1000); 10 min; All at RT |
